# Retinal circuits driving a non-image forming visual behavior

**DOI:** 10.1101/2020.09.08.288373

**Authors:** Corinne Beier, Ulisse Bocchero, Zhijing Zhang, Nange Jin, Stephen C. Massey, Christophe P. Ribelayga, Kirill Martemyanov, Samer Hattar, Johan Pahlberg

**Author notes:** These authors contributed equally to this work (Corinne Beier, Ulisse Bocchero). Co-correspondence, Editorial Correspondence: Samer Hattar, PhD, Section on Light and Circadian Rhythms, National Institute of Mental Health (NIMH), National Institutes of Health (NIH), Building 35A, Room 2E-450, Bethesda, MD 20892, USA. **Author Contributions**, C.B., U.B, K.M., S.H., and J.P., conceptualized the project and designed the methodology. C.B. performed the investigation and data analysis of PLR experiments. U.B. performed the electrophysiology experiments, U.B. and J.P. analyzed the electrophysiology data. Z.Z, N.J., S.C.M, and C.P.R. performed the investigation and data analysis of the cone-Cx36 KO experiments. K.M. provided resources for the experiments. C.B. contributed to writing – original draft preparation. C.B., U.B, K.M, S.H., J.P., contributed to writing – review and editing.

## Abstract

Outer retinal circuits that drive non-image forming vision in mammals are unknown. Rods and cones signal light increments and decrements to the brain through the ON and OFF pathways, respectively. Although their contribution to image-forming vision is known, the contributions of the ON and OFF pathway to the pupillary light response (PLR), a non-image forming behavior, are unexplored. Here we use genetically modified mouse lines, to comprehensively define the outer retinal circuits driving the PLR. The OFF pathway, which mirrors the ON pathway in image-forming vision, plays no role in the PLR. We found that rods use the primary rod pathway to drive the PLR at scotopic light levels. At photopic light levels, the primary and secondary rod pathways drive normal PLR. Importantly, we find that cones are unable to compensate for rods. Thus, retinal circuit dynamics allow rods to drive the PLR across a wide range of light intensities.

## Introduction

Animals perceive their visual environment, both consciously and subconsciously, when the retina conveys light information to image forming and non-image forming vision. In the mammalian retina, there are three types of photoreceptors, the outer retinal photoreceptors, rods and cones, and intrinsically photosensitive retinal ganglion cells (ipRGCs). It has been demonstrated that outer retinal photoreceptors and ipRGCs contribute to the pupillary light response (PLR), a non-image forming behavior (Butler and Silver, 2011; Hattar et al., 2003; Kostic et al., 2016; Lall et al., 2010; Lucas et al., 2001, 2003; Panda et al., 2003). The melanopsin (Opn4) contribution to pupil constriction, the photopigment behind the intrinsic light response in ipRGCs, is limited to high light intensity (Keenan et al., 2016; Lucas et al., 2003). The rod and cone contribution to pupil constriction is still controversial, but more recent research suggests that rods are the predominant driver of the PLR (Keenan et al., 2016). Surprisingly however, even elemental aspects of the outer retinal circuits behind the photoreceptor-mediated PLR are still unexplored.

In image forming vision, there are two well-documented parallel pathways, the ON and OFF pathways, which respond to the onset or offset of light, respectively (reviewed by Demb and Singer, 2015). This split in information along parallel pathways begins at the first synapse in the retina, where rod and cone photoreceptors synapse with ON and OFF bipolar cells, which express mGluR6 and ionotropic glutamate receptors, respectively. Except at high light levels, rods are thought to be the predominant contributor to pupil constriction across a range of light intensities (Keenan et al., 2016). There are three rod pathways in the retina that are known to contribute to image forming vision: the primary, secondary and tertiary rod pathways (Figure 1A) (reviewed in Demb and Singer, 2012; Grimes et al., 2018; Völgyi et al., 2004). In the most sensitive pathway (Dacheux and Raviola, 1986; Grimes et al., 2018b; Ke et al., 2014), the primary rod pathway, rods are synaptically connected to rod bipolar cells. Rod bipolar cells synapse with AII amacrine cells which convey the rod signal to the cone pathway via a gap junction to ON-cone bipolar cells and to OFF-cone bipolar cells via an inhibitory glycinergic synapse (Figure 1A, green) (Dacheux and Raviola, 1986). In the secondary rod pathway, rod signals are routed to cones via rod/cone gap junctions and then conveyed to ON and OFF cone bipolar cells (Figure 1A, purple) (Dacheux and Raviola, 1982; Nelson, 1977). In the tertiary pathway, rods are thought to synapse directly with OFF-cone bipolar cells (Figure 1A, blue) (Li et al., 2004). Although these pathways have been well-studied, it remains unknown if and how the PLR relies on these same pathways.

**Figure 1.**
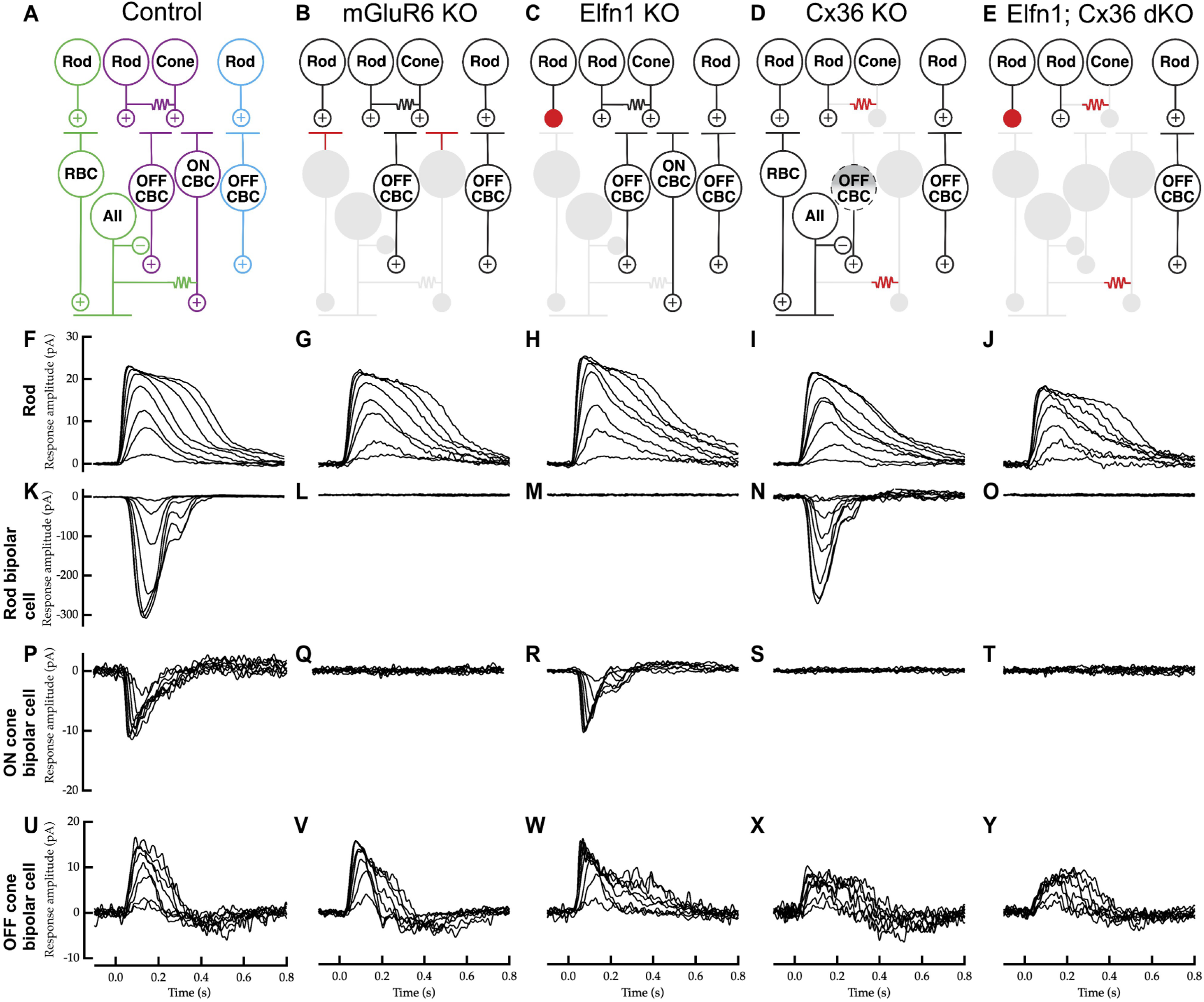
Rod circuit pathways in control and mutant mice. (**A**) A schematic of the three rod pathways in the mammalian outer retina. In the primary rod pathway (green) the rod signal is passed from rods to rod bipolar cells (RBCs) to AII amacrine cells. AII amacrines convey the signal via a gap junction connection to ON-cone bipolar cells (ON CBCs) and via an inhibitory glycinergic synapse to OFF-cone bipolar cells (OFF CBCs). In the secondary rod pathway (purple) the rod signal is passed from rods to cones via a gap junction connection and then from cones to both ON and OFF CBCs. In the tertiary rod pathway (blue) the rod signal is passed from rods to OFF CBCs. (**B**) A retinal circuit schematic of the mGluR6 knockout (KO) mouse showing functional and non-functional rod pathways. RBCs and ON CBCs cannot receive glutamate signal due to the loss of the metabotropic glutamate receptor 6 (mGluR6) at their dendrites (red). The downstream rod ON pathway no longer receives rod input (gray). (**C**) A retinal circuit schematic of the Elfn1 KO mouse showing functional and non-functional rod pathways. Due to the loss of the presynaptic protein Elfn1, the synapse between rods and RBCs is non-functional (red). The primary pathway no longer receives rod input (gray). (**D**) A retinal circuit schematic of the Cx36 KO mouse showing functional and non-functional rod pathways. Due to the loss of connexin 36 (Cx36) no gap junctions are formed between rods and cones, as well as between AII and ON CBCs (red). The secondary pathway no longer receives rod input (gray). OFF CBCs can still receive inhibitory scotopic light input from the primary rod pathway, but not excitatory scotopic input from the secondary rod pathway (indicated by gray gradient). (**E**) A retinal circuit schematic of the Elfn1; Cx36 double KO (dKO) mouse showing functional and non-functional rod pathways. The synapse between the rod and RBC is non-functional and there is a lack of rod to cone and AII to ON-CBCs gap junctional coupling, respectively (red). The primary and secondary rod pathways no longer receive rod input (gray). (**F-J**) Physiological recordings of rod photocurrents in retinal slices from control, mGluR6 KO, Elfn1 KO, Cx36 KO and Elfn1; Cx36 dKO mice, respectively. Recordings were made in wholecell patch clamp mode (V_m_ = −40 mV). 20 ms light flashes were given at time 0 s. Flash strengths range from 2.5 to 156 R*/rod. Recordings are representative of data collected across several cells (see Supplementary Table 1). (**K-O**) Physiological recordings of light-evoked RBC responses from control, mGluR6 KO, Elfn1 KO, Cx36 KO and Elfn1; Cx36 dKO mice, respectively. Recordings were made in wholecell patch clamp mode (V_m_ = −60 mV). Flash strengths range from 0.2 to 16 R*/rod. RBC light responses were never observed in mGluR6 KO, Elfn1 KO or Elfn1; Cx36 dKO mice (Supplementary Table 1). Recordings were from the same slices as the rod recordings. (**P-T**) Physiological recordings of light-evoked ON CBC responses as described for RBCs. Flash strengths spanned a range expected to activate the primary and secondary rod pathways. Recordings were from the same slices as the rod and RBC recordings, typically no more than 50 μm apart from each other. ON CBC light responses were never observed in mGluR6 KO, Cx36 KO or Elfn1; Cx36 dKO mice (Supplementary Table 1). (**U-Y**) Physiological recordings of light-evoked OFF CBCs responses in retinal slices as described above. Flash strengths spanned a range expected to activate the primary, secondary and tertiary rod pathways. Recordings were from the same slices as the rod, RBC and/or ON CBC recordings, typically within 50 μm from each other (Supplementary Table 1).

In this study, we used a multitude of genetically modified mice to systematically and precisely dissect the rod and cone pathways in the retina to investigate their contributions to a non-image forming behavior. We show, with electrophysiological evidence, that the genetic mutant mouse lines dissect the rod and cone circuits, allowing us to investigate their contribution to the PLR across a range of light intensities. Specifically, we find that the ON pathway is required for the PLR, and the OFF pathway plays no role, in stark contrast to image forming vision. Furthermore, we show, for the first time, that a non-image forming visual behavior requires the most sensitive retinal pathway, the primary rod pathway, for normal function. This study reveals the rod circuits that drive the PLR at all light intensities.

## Results

### The ON pathway is required for the photoreceptor-mediated PLR

We began our investigation by asking if the PLR requires both major parallel pathways in the retina, the ON and OFF pathways. At the first synapse in the retina, light information is sent to ON and OFF bipolar cells, which respond to the onset or offset of light, respectively. In the mammalian retina, synaptic transmission to the ON bipolar cells is mediated by the postsynaptic metabotropic glutamate receptor 6 (mGluR6) (Masu et al., 1995; Nakajima et al., 1993; Nomura et al., 1994). To isolate the OFF pathway, we silenced the ON pathway using the mGluR6 knockout (KO) animals (Maddox et al., 2008). In these animals, the primary rod pathway and ON-cone bipolar cells are unresponsive to light. The rod secondary OFF, rod tertiary, and cone OFF pathways are functional (Figure 1B).

To confirm the behavior of single retinal neurons in a network, we performed whole-cell patch clamp recordings from dark-adapted retinal slices. Rods in control and mGluR6 KO mice showed robust light evoked responses, and our measurements agree well with responses seen in the literature (Figure 1F and 1G) (Pahlberg et al., 2017). After confirming that rods in our mice were functional, we recorded light responses from the immediate downstream neurons in the retina (Figure 1A); rod bipolar cells (RBCs), ON cone bipolar cells (ON CBCs), and OFF cone bipolar cells (OFF CBCs). In control mice, RBCs and ON CBCs show robust depolarizing light responses, as expected (Figure 1K and 1P) (Cao et al., 2012; Field and Rieke, 2002; Okawa et al., 2010). In mGluR6 KO mice, the light response in both RBCs and ON CBCs were totally absent, as predicted (Figure 1L and 1Q) (Xu et al., 2012). In both control mice and mGluR6 KO mice OFF CBCs showed robust hyperpolarizing responses with similar sensitivity (Figure 1U and 1V, Supplementary Table 1). These results confirm the ON pathway is silenced in our mGluR6 KO mice.

We investigated how the mGluR6 loss of function affects the PLR by observing dark-adapted pupil constriction to overhead light. Light intensity was set to a photopic (100 lux) light level and kept on for the entire recording period (see Methods). In the photopic regime, mGluR6 KO mice show impaired pupil constriction compared to controls (Figure 2B). Interestingly, these animals exhibited a delayed response (Figure 2A) and showed a deficit in constriction immediately following light onset compared to controls (ANOVA post hoc Dunnett’s test, p = 0.002). We quantified this further by measuring the time to half the minimum pupil constriction and found that mGluR6 KO mice are delayed compared to controls (Controls 1.5 ± 0.2 seconds, mGluR6 KO 4.9 ± 1.2 seconds to half constriction, ANOVA post hoc Dunnett’s test, p = 0.004). The delay we observe in the PLR resembles the response seen in animals where ipRGCs are the only functioning photoreceptor (Keenan et al., 2016; Lucas et al., 2001). To determine if the remaining response in mGluR6 KO animals is entirely due to ipRGCs and to isolate any remaining photoreceptor-mediated PLR, we generated mGluR6; Opn4 double knockout (dKO) animals. These animals allowed us to silence the ON pathway and the melanopsin (Opn4) response at the same time, while leaving the OFF pathway completely functional (Figure 1B and 1V). The PLR is entirely abolished in mGluR6; Opn4 dKO mice (Figure 2A and 2B), while Opn4 KO animals show only a small, non-significant, deficit (Keenan et al., 2016; Lucas et al., 2003) (data not shown). Therefore, at photopic light levels, the OFF pathway alone cannot contribute to the PLR.

**Figure 2.**
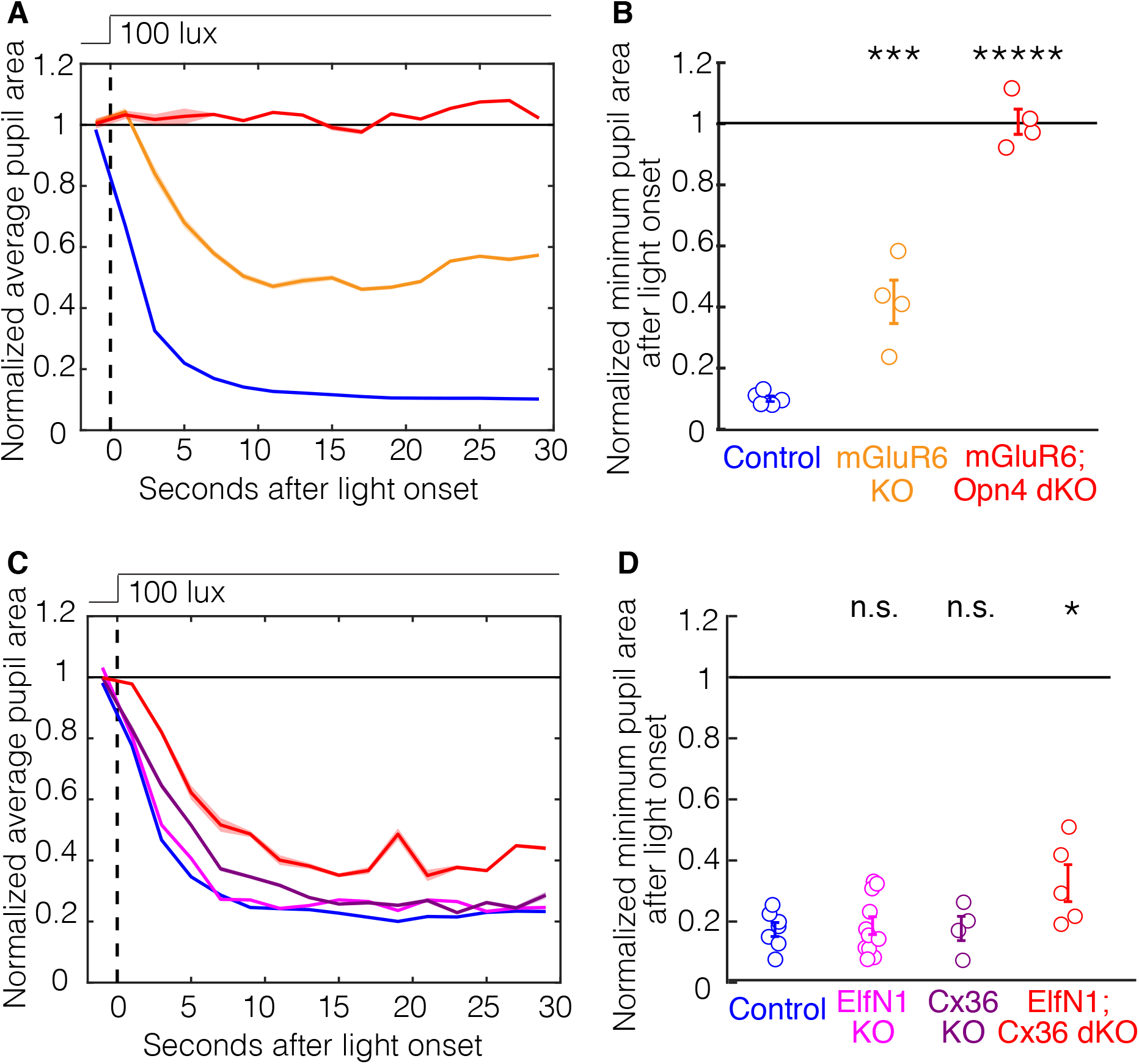
The rod ON pathways drive the photopic pupillary light response. (**A**) The average pupil constriction over time in response to 100 lux light beginning at t = 0 seconds (dashed line) for control (blue), mGluR6 KO (orange) and mGluR6; Opn4 double KO (red) mice. Shaded outlines represent SEM. All pupil sizes are normalized to the dark-adapted pupil size (before t = 0). mGluR6 KO mice (orange) have a pupil constriction deficit immediately following light onset compared to control mice (ANOVA post hoc Dunnett’s method, p = 0.002). (**B**) The minimum pupil area (maximal constriction) in response to 100 lux light from t = 0 to t = 30 seconds. All pupil sizes are normalized to the dark-adapted pupil size (before t = 0) Individuals are shown as circles. Error bars show standard error on the mean (SEM). Significance from control group is as follows: mGluR6 KO p = 8E-4, mGluR6; Opn4 dKO p = 9E-8 (ANOVA post hoc Tukey-Kramer method). (**C**) The average pupil constriction over time in response to 100 lux light beginning at t = 0 seconds (dashed line) for control (blue), Elfn1 KO (magenta), Cx36 KO (dark purple), and Elfn1; Cx36 dKO (red) mice. Shaded outlines represent SEM. All pupil sizes are normalized to the dark-adapted pupil size (before t = 0). Elfn1; Cx36 dKO mice (red) have a pupil constriction deficit immediately following light onset compared to control mice (ANOVA post hoc Dunnett’s method, p = 0.04). (**D**) The minimum pupil area (maximal constriction) in response to 100 lux light from t = 0 to t = 30 seconds. Individuals are shown as circles. Error bars show SEM. Significance from control group is as follows: Elfn1 KO p = 0.992, Cx36 KO p = 0.996, Elfn1; Cx36 dKO p = 0.0498 (ANOVA post hoc Tukey-Kramer method).

### Both the primary and secondary rod pathways drive the photopic PLR

We next silenced the primary rod pathway by knocking out the pre-synaptic cell-adhesion molecule Elfn1, that specifically prevents the formation of the rod to RBCs synapse (Figure 1C) (Cao et al., 2015). We again recorded light evoked responses in single cells from dark adapted retinal slices. Rod responses in control and Elfn1 KO mice were normal (Figure 1H). However, RBCs in Elfn1 KO mice did not respond to light flashes, indicating a lack of synaptic contact and signaling from rods to RBCs (Figure 1M) (Cao et al., 2015). By contrast, ON and OFF CBCs responses in Elfn1 KO mice were completely normal (Figure 1R and 1W). Therefore, the Elfn1 mutation specifically silences signaling in the primary rod pathway (Cao et al., 2015). Using photopic light intensities, we observe Elfn1 KO mice have no PLR deficits (Figure 3C and 3D), indicating that the primary rod pathway is dispensable for normal photopic PLR.

**Figure 3.**
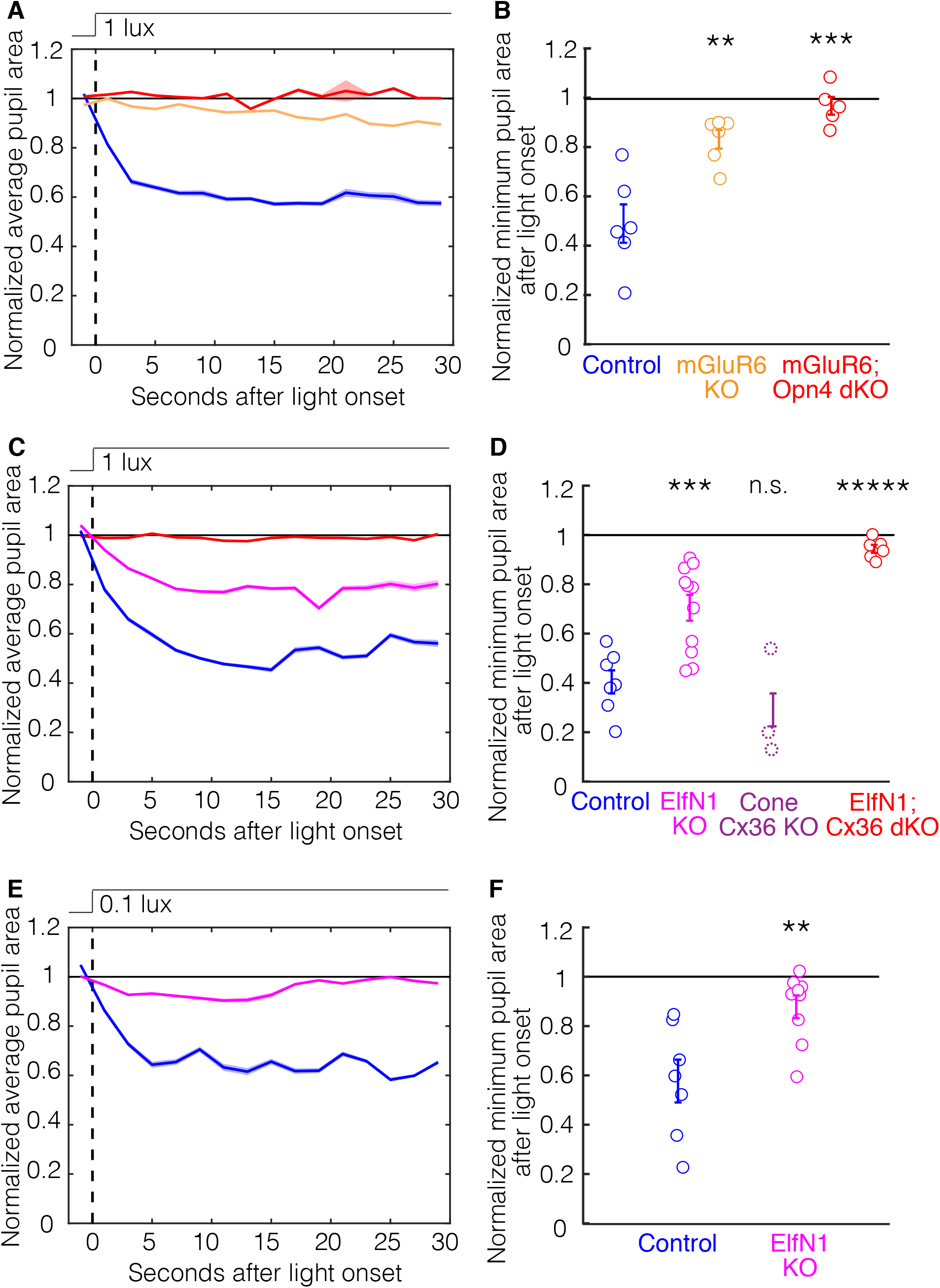
The primary rod pathway is required for the scotopic pupillary light response. (**A**) The average pupil constriction over time in response to 1 lux light beginning at t = 0 seconds (dashed line) for control (blue), mGluR6 KO (orange) and mGluR6; Opn4 double KO (red) mice. Shaded outlines represent SEM. All pupil sizes are normalized to the dark-adapted pupil size (before t = 0). (**B**) The minimum pupil area (maximal constriction) in response to 1 lux light from t = 0 to t = 30 seconds. Individuals are shown as circles. Error bars show standard error on the mean (SEM). Significance from control group is as follows: mGluR6 KO p = 0.007, mGluR6; Opn4 dKO p = 3E-4 (ANOVA post hoc Tukey-Kramer method). (**C**) The average pupil constriction in response to 1 lux light beginning at t = 0 seconds (dashed line) for control (blue), Elfn1 KO (magenta), and Elfn1; Cx36 double KO (red) mice. Shaded outlines represent SEM. All pupil sizes are normalized to the dark-adapted pupil size (before t = 0). (**D**) The minimum pupil area (maximal constriction) in response to 1 lux light from t = 0 to t = 30 seconds. Individuals are shown as circles. Error bars show SEM. Significance from control group is as follows: Elfn1 KO p = 5E-4, Elfn1; Cx36 dKO p = 1E-6 (ANOVA post hoc Tukey-Kramer method). Cone-Cx36 KO mice are indicated in purple dashed circles. These mice were not significant from control mice shown in Supplementary Figure 1B (Student’s paired t test, p = 0.65) (**E**) The average pupil constriction over time in response to 0.1 lux light beginning at t = 0 seconds (dashed line) for control (blue) and Elfn1 KO (magenta) mice. Shaded outlines represent SEM. All pupil sizes are normalized to the dark-adapted pupil size (before t = 0). (**F**) The minimum pupil area (maximal constriction) in response to 0.1 lux light from t = 0 to t = 30 seconds. Individuals are shown as circles. Error bars show SEM. Significance of Elfn1 KO mice from control group is p = 0.006 (Student’s two-way t-test).

To investigate whether the secondary rod pathway is required at photopic light levels for normal PLR we used connexin 36 (Cx36) KO mice (Deans et al., 2001). Rod signals are passed to cones via a Cx36 gap junction and from AIIs to ON-cone bipolar cells (Figure 1D) (Deans et al., 2002; Güldenagel et al., 2001). When recording from Cx36 KO mice we again found that, as with the other two mutations, rods show robust normal responses (Figure 1I). Importantly, RBC responses in these mice were normal (Figure 1N). In total agreement with the Cx36 mutation, ON CBCs do not show responses, whereas OFF CBCs show responses with normal sensitivity (Figure 1S and 1X) (Deans et al., 2002; Güldenagel et al., 2001).

Remarkably, Cx36 KO mice show normal photopic PLR with the same kinetics as control mice. These results imply that either the rod to rod bipolar cell pathway, independent of the AII ON pathway, is capable of driving pupil constriction or that cones are playing a role. If rods are the only outer retinal photoreceptor driving the photopic PLR, however, we predict that silencing the primary and secondary rod pathways together will give a melanopsin only PLR.

To completely silence the primary and secondary rod pathways we generated Elfn1; Cx36 dKO mice (Figure 1E). Importantly, the cone pathways are unaffected in these mice. After confirming that rod light responses in Elfn1; Cx36 dKO in retinal slices are comparable to controls (Figure 1J), we assessed the downstream effects. As predicted, the light evoked responses in both RBCs and ON CBCs were absent (Figure 1O and 1T). Interestingly, we were able to record OFF CBC light responses in these mice, directly showing for the first time the isolated tertiary rod pathway light response, without any input from the primary or secondary rod pathways (Figure 1Y) (Supplementary Table 1) (Pang et al., 2012; Tsukamoto et al., 2001). The sensitivity in the Elfn1; Cx36 dKO OFF CBCs were very similar to controls, indicating that the direct rod input in the tertiary pathway can drive OFF CBCs near normal levels in the absence of the secondary pathway (Supplementary Table 1).

Fascinatingly, when we observe the PLR of Elfn1; Cx36 dKO mice we see a delayed pupil response to light onset (ANOVA post hoc Dunnett’s test, p = 0.04) in addition to a small pupil constriction deficit (Figure 2C). We again quantified the delay by measuring the time to half the minimum pupil constriction. We find that Elfn1; Cx36 dKO mice show a delayed PLR compared to controls (Controls 2.6 ± 0.5 seconds and mGluR6 KO 5.9 ± 2.0 seconds to half constriction, ANOVA post hoc Dunnett’s test, p = 0.03). The delayed response resembles the melanopsin-dependent PLR we see in our mGluR6 KO animals (Figure 2A), indicating that cones cannot compensate for the rod contribution to the photopic PLR.

In conclusion, the primary and secondary rod pathways can compensate for one another to drive a normal photopic PLR, but the loss of both pathways results in PLR deficiencies. We next investigated the role these circuits play in the scotopic PLR.

### The primary rod pathway has a differential effect on the scotopic PLR

To study the contribution of the ON pathway to the scotopic PLR, we measured the pupil constriction of mGluR6 KO mice in response to dim light exposure (1 lux). We found that mGluR6 KO mice have a severe PLR deficit compared to controls but were surprised to see a very weak and slow pupil constriction (Figure 3A and 3B, one-way Student’s t-test from dilation, p = 0.007). This slow response is indicative of melanopsin and it disappeared in mGluR6; Opn4 dKO mice, which have no scotopic PLR (Figure 3A and 3B, one-way Student’s t-test from dilation, p = 0.4). These results demonstrate that the ON pathway is required for the scotopic PLR and that the rod tertiary pathway plays no role.

We then examined the scotopic PLR of Elfn1 KO mice to investigate the primary rod pathway contribution to pupil constriction in low light. Although we find a large deficit in the Elfn1 KO mouse PLR it is clear that there is some pupil constriction (Figure 3C and 3D). To investigate the role of the secondary rod pathway to the PLR in low light, we used a Cone-Cx36 KO mouse, to eliminate the connexin only in cone cells leading to the abolition of electrical synapses between rods and cones (Jin et al., 2020). The conditional cone specific connexin knockout, importantly, leaves the AII Cx36 connections in the primary rod pathway intact (compare Figure 1D to Supplementary Figure 1A). Cone-Cx36 KO mice had an indistinguishable pupil constriction at 1 lux compared to controls (Figure 3D and Supplementary Figure 1). To determine if the remaining response in the ElfN1 KO mice is due to the secondary pathway, we looked at the scotopic PLR in Elfn1; Cx36 dKO mice. Silencing the primary and secondary rod pathways abolishes the scotopic PLR (Figure 3C and 3D), which is more severe than mGluR6 KO mice (see Discussion). Therefore, although dispensable at photopic light levels, we find that the primary rod pathway is required for normal PLR at scotopic light levels and that the secondary rod pathway cannot compensate for its loss. This is the first proof that non-image forming vision requires the most sensitive retinal pathway for normal function.

For image forming vision, the smallest photon capture events are thought to be conveyed by the primary rod pathway, which is more sensitive than the secondary and tertiary rod pathways (Supplementary Table 1) (Grimes et al., 2014; Ke et al., 2014). To investigate if this difference in pathway contribution holds true for non-image forming vision, we looked at the PLR of Elfn1 KO mice at an even lower light level (0.1 lux). Under these conditions Elfn1 KO mice have almost no PLR (Figure 3E and 3F). This result indicates that the primary rod pathway is the main driver behind the PLR at very low light levels.

## Discussion

Here, we define the outer retinal circuits driving the PLR at scotopic and photopic light intensities. Remarkably, we find that rod image-forming visual pathways play distinct roles in the PLR. Specifically, the primary and secondary rod pathways contribute to the photopic PLR, with no contribution from cones. We show that non-image forming vision, similar to image vision, requires the most sensitive retinal pathway for normal function at scotopic light levels.

### The OFF pathway alone does not contribute to the PLR

Mice with only a functional OFF pathway are capable of driving image-forming vision (Iwakabel et al., 1997). In fact, these mice show completely normal OFF responses (Figure 1Y). However, the OFF pathway alone does not drive pupil constriction in response to light at both photopic and scotopic light levels. It is clear that the ON and OFF pathways do not play the same role in non-image forming vision.

In the original mGluR6 KO mouse studies (Iwakabel et al., 1997), the authors suggested that the OFF pathway contributes to the PLR. However, this study occurred before the discovery of ipRGCs. Here, we show that the remaining pupil constriction in mGluR6 KO mice is due to melanopsin by two ways. First, the mGluR6 KO mice showed a delayed pupil constriction similar to the delay observed in animals lacking rods and cones (Lucas et al., 2001). Second, in the mGluR6; Opn4 dKO mice, all pupil constriction disappears showing that the response is entirely due to melanopsin.

### Cones are unable to drive the PLR, even at photopic light levels

Mice lacking either the primary or secondary rod pathway have no pupil constriction deficits at photopic light levels. In addition, our mutant lines show that either pathway is sufficient to drive the photopic PLR. Specifically, mice with both the primary and secondary rod pathways silenced showed a delay in the photopic PLR kinetics. In these mice, cone pathways are completely functional, therefore, the fast kinetics of the PLR in the single Elfn1 KO and Cx36 KO mice can only be attributed to rods (Cao et al., 2015; Deans et al., 2002), consistent with data from Gnat1; Opn4 dKO mice, where cones are the only functional photoreceptors (Keenan et al., 2016). Furthermore, the PLR in the Elfn1; Cx36 dKO mice resembles the PLR we observe in mGluR6 KO mice, which was entirely due melanopsin. This shows that cones play little to no role in the PLR.

### Non-image forming vision requires the most sensitive retinal pathway for normal function

Electrophysiological evidence suggested that ipRGCs receive light information from the primary rod pathway, but it remains unknown if this connection is behaviorally relevant (Weng et al., 2013). We tested this directly by using mice lacking the primary rod pathway and showed a clear pupil constriction deficit at low light levels. The contribution of the most sensitive rod pathway becomes even clearer at lower light levels when Elfn1 KO mice have almost no pupil constriction. Mice lacking the secondary rod pathway have no pupil constriction deficits at low light, when the primary pathway is fully intact as we have demonstrated in the conditional connexin 36 knockout. This shows that the primary rod pathway fully compensates for the secondary pathway at scotopic light levels, but importantly, the secondary pathway does not fully compensate for the primary pathway. Our results, therefore, show that the primary pathway is the predominate contributor to the scotopic PLR (Figure 4B).

**Figure 4.**
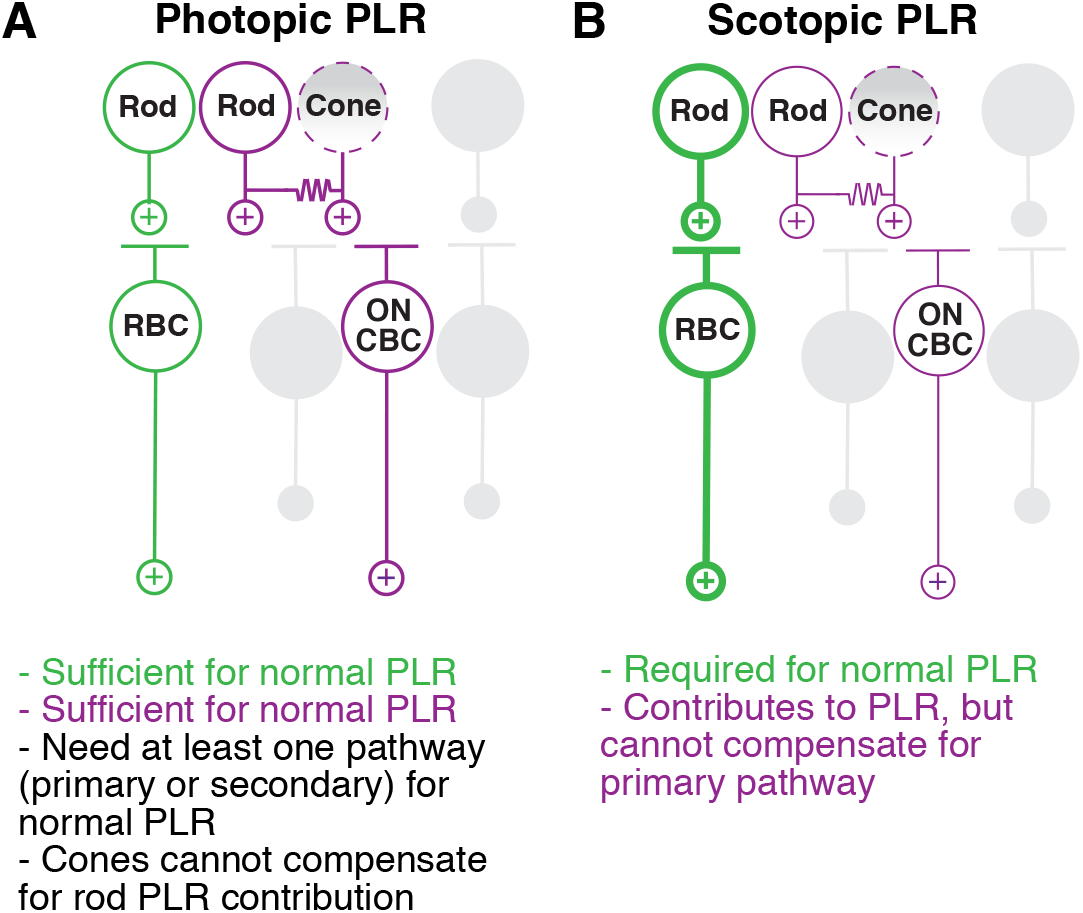
The outer retinal circuit pathways driving the pupillary light response. (**A**) A schematic detailing the outer retinal circuit pathways that drive the photopic PLR. The primary rod pathway (green) and the secondary ON rod pathway (purple) can drive normal PLR. The secondary OFF rod pathway and tertiary pathway play no role (gray). Cones cannot compensate for rods even when the secondary rod pathway drives the photopic PLR (indicated by dashed outline and shaded gradient). The potential role of the AII amacrine cell (not shown in schematic) is discussed in the manuscript and Supplementary Figure 1. (**B**) A schematic detailing the outer retinal circuit pathways that drive the scotopic PLR. The primary rod pathway is the predominate circuit driving the scotopic PLR (bolded green) and is required for normal pupil constriction. The secondary ON rod pathway (purple) can contribute to pupil constriction but cannot compensate for the primary rod pathway for normal PLR. The secondary OFF rod pathway and tertiary rod pathway play no role (gray). Cones cannot compensate for rods at scotopic light levels (indicated by dashed outline and shaded gradient). The potential role of the AII amacrine cell (not shown in schematic) is discussed in the manuscript and Supplementary Figure 1.

### A dramatically slow PLR at scotopic levels depends on the secondary rod OFF pathway

Melanopsin has long been thought to be relatively insensitive to light (but see Lee et al., 2019) and cannot induce pupil constriction when it is the only functioning photoreceptor except in very bright light (Keenan et al., 2016; Lucas et al., 2001). Surprisingly, we find that melanopsin drives an incredibly slow pupil constriction at scotopic light levels in mGluR6 KO animals (Figure 3A). The slow pupil constriction disappears in mGluR6; Opn4 dKO animals. The Elfn1; Cx36 dKO mice, which also contain melanopsin, fail to show this slow response (Figure 3C). The only difference between mGluR6 KO and Elfn1; Cx36 dKO mice is a functioning secondary OFF rod pathway (Figure 1B and 1E). This suggests that melanopsin can contribute to scotopic pupil constriction, but only in combination with the rod-cone to OFF cone bipolar cell pathway. We hypothesize that the hyperpolarization of the OFF pathway, in response to light onset, can increase the melanopsin-mediated ipRGC sensitivity, and therefore contribution to the PLR (Wong et al., 2007).

### The primary and secondary rod pathways drive the photopic PLR

Our results showing that both the OFF pathway and cones are not capable of driving the PLR, leaves the primary and secondary ON rod pathways as possible circuits behind the photoreceptor-mediated photopic PLR. Mice lacking the primary rod pathway retain a normal photopic PLR, indicating that the secondary ON rod pathway is sufficient to drive normal pupil constriction in photopic light (Figure 4A). When the secondary pathway is silenced, the PLR is also normal. Therefore, the rod to rod bipolar cell synapse is sufficient for the fast photoreceptor-mediated PLR (Figure 4A).

How do the rod bipolar cells convey the signal to downstream circuitry to drive the photopic PLR? We were surprised to find that Cx36 KO mice, which lack the connection between AII amacrine cells and ON cone bipolar cells of the primary rod pathway, still show normal PLR at photopic light levels. This data leads to a possible explanation that the rod bipolar cells synapse directly onto the output neurons of the retina to drive the PLR (Supplementary Figure 2A). It has been well documented that ipRGCs drive pupil constriction in response to light (Chen et al., 2011; Güler et al., 2008; Hatori et al., 2008). Specifically, one type of ipRGC, the M1 ipRGC, is required for normal PLR and there is anatomical evidence that suggests a direct rod bipolar cell input to M1 ipRGCs (Østergaard et al., 2007).

Another possible circuit mechanism to drive the photopic PLR from the rod bipolar cells is via the AII inhibitory glycinergic synapse. In Cx36 KO mice, the AII to OFF cone bipolar cell pathway is still active (Figure 1D) (Graydon et al., 2018). However, to-date there is no anatomical evidence showing direct OFF cone bipolar cell to ipRGC synaptic connections (Sabbah et al., 2018). Intriguingly, the AII inhibitory synapses are found at the same stratification layer within the retina as M1 ipRGC dendrites (Sabbah et al., 2018; Tsukamoto and Omi, 2017). We propose an indirect circuit between AII inhibitory synapses and M1 ipRGCs, likely via another amacrine cell such as the dopaminergic amacrine cell, to disinhibit the photopic PLR (Supplementary Figure 2B) (Newkirk et al., 2013; Pérez-Fernández et al., 2019; Roy and Field, 2019).

There is a third possible pathway that may drive the PLR from the rod to rod bipolar cell synapse, although we show it is not required for the photopic PLR. In the primary rod pathway, AII amacrine cells make gap junction connections with multiple ON cone bipolar cell types. The AII favors connections with Type 6 ON cone bipolar cells (Tsukamoto and Omi, 2017), which synapse with most ipRGC subtypes and is the primary bipolar cell partner to M1 ipRGCs, making this downstream pathway a possible contender driving the PLR (Supplementary Figure 2C) (Dumitrescu et al., 2009; Sabbah et al., 2018).

## Methods

### Animals

All mice were handled in accordance with guidelines of the Animal Care and Use Committees of the National Institute of Mental Health (NIMH). Male and female mice, aged 2 to 6 months were used in experiments. mGluR6 KO were obtained from The Jackson Laboratory (Stock number 016883) (Maddox et al., 2008) and crossed with Opn4 knockout mice (Ecker et al., 2010) to obtain control mice (mGluR6+/-; Opn4 +/-) and double knockout mice (mGluR6-/-; Opn4 -/-), in addition to mGluR6 KO littermates (mGluR6-/-; Opn4 +/-). Elfn1 KO mice (Cao et al., 2015) were crossed with Cx36 KO mice (a gift from the Dr. Jeffrey Diamond laboratory) (Deans et al., 2001) to generate control mice (Elfn1 +/+; Cx36 +/+), Elfn1 KO mice (Elfn1-/-; Cx36 +/+), Cx36 KO mice (Elfn1 +/+; Cx36 -/-), and Elfn1; Cx36 double knockout mice (Elfn1 -/-; Cx36 -/-). Mice were group housed in a temperature and humidity controlled room under a 12 hour light-dark cycle with light intensity near 100 lux. Food and water were available *ad libitum*.

Cone-Cx36 KO mice are described in (Jin et al., 2020).

### Single-cell electrophysiological recordings

Light-evoked responses were made from dark-adapted retinal slices as previously described (Pahlberg et al. 2017). Briefly, mice were dark adapted overnight and euthanized according to protocols and guidelines approved by NIH. Their eyes were enucleated and hemisected under infrared light. Retinas were extracted from the eye cups, embedded in low density agar gel (3%) and cut with a vibratome to obtain 200 μm thick slices. Retina slices were perfused with Ames’ medium with an 8 ml/min flow rate (equilibrated with 5% CO_2_/ 95% O_2_), maintained at 35–37 °C, and visualized in the infrared. The pipette internal solution for wholecell patch clamp recordings contained (in mM): 125 K-aspartate, 10 KCl, 10 HEPES, 5 N-methyl glucamine-HEDTA, 0.5 CaCl2, 1 ATP-Mg and 0.2 GTP-Mg; pH was adjusted to 7.4 with NMG-OH. Light responses were generated delivering 20 ms flashes from a blue-green light emitting diode (λ_max_ ~ 505 nm). Flash strengths were varied from a just-measurable response to those that produced a maximal response, increasing in factors of two. Membrane currents were filtered at 300 Hz and sampled at 10 kHz. Light sensitivity for each cell type was estimated from the half-maximal flash strength from the best Hill fit, for each genotype. To identify RBCs, ON CBCs, and OFF CBCs the shape and morphology of the cells, the time course, the amplitude and the polarity of their light responses were considered. As a control, the electrode internal solution for some experiments contained Alexa-750 (100 μM), which allowed visualization of the cells in the far red without significantly bleaching the visual pigment (see Supplementary Figure 3). The subtypes of ON and OFF CBCs were not characterized. All recordings were performed between circadian time 4 and 12.

### Pupillometry

The pupillary light response (PLR) experiments were performed as previously described (Keenan et al., 2016). Briefly, mice were handled prior to PLR experiment days to habituate animals to handling. On the day of the experiment, mice were dark adapted for at least 1 hour before being scruffed and placed in front of a fixed-focus video camera (Sony 4K HD Camcorder FDRAX3) under infrared light (ICAMI IR Illuminators, λ = 850 nm). The dark-adapted pupils were recorded for at least 5 seconds before turning the stimulus light on. The stimulus light bulb (Sunlite 6500K A19/LED/10W) was placed directly above the mouse such that the stimulus provided 100 lux light as measured by a lux meter (EXTECH Foot Candle/Lux Light Meter, 401025). For experiments under scotopic conditions, neutral density filters were placed over the light to achieve 1 lux and 0.1 lux stimuli. Pupils were recorded for at least 30 seconds following light onset. All recordings were performed between Zeitgeber times 4 and 12.

A semi-automated custom script was used to measure the pupil area in every video frame (30 frames/second). For each mouse, all measurements were normalized to the average dark-adapted pupil size across the 5 seconds immediately prior to light onset. Minimum pupil area (maximum pupil constriction) was taken from pupil measurements between t = 0 to 30 seconds after light onset.

Pupillometry experiments for Cone-Cx36 KO mice and their controls were performed as above except that pupil size was recorded at t = 0, 5, 15, and 30 seconds. Minimum pupil area measurements were taken from t = 30.

### Statistical analysis

Statistical analyses are listed in the text and Figure legends. Number of animals used are indicated in the Figures and Supplementary Table 1.

## Acknowledgements

This project was supported in part by the funding from the Intramural Research Programs of the NIDCR, NINDS and NEI at the National Institutes of Health (J.P.) and from funding from the National Institute of Mental Health project number MH002964 (S.H.). We would like to thank Hui Wang in the Samer Hattar lab for her management of the Opn4^Cre^ mouse line. We would like to also thank Hua Tian in the Jeff Diamond lab for her management of the Cx36 knockout line shared with us.

## Supplementary Figure Legends

**Supplementary Figure 1. The secondary rod pathway is not required for the scotopic pupillary light response**

(**A**) A retinal circuit schematic of the Cone-Cx36 KO mouse showing functional and non-functional rod pathways. Due to the loss of connexin 36 (Cx36) in cones, no gap junctions are formed between rods and cones (red). The secondary pathway no longer receives rod input (gray). ON and OFF CBCs can still receive excitatory and inhibitory scotopic light input from the primary rod pathway, respectively (indicated by gray gradients).

(**B**) The minimum pupil area (maximal constriction) in response to 1 lux light from t = 0 to t = 30 seconds. Individuals are shown as circles. Error bars show SEM. Cone-Cx36 KO mice are not significantly different from controls (Student’s paired t test, p = 0.65)

**Supplementary Figure 2. Possible rod bipolar cell to ipRGC circuits driving the PLR.**

The rod to RBC synapse can drive normal photopic and scotopic PLR. We propose the following circuits to describe how the RBC signal might be conveyed to ipRGCs to induce pupil constriction.

(**A**) A schematic showing RBCs making direct synapses with the M1 ipRGC subtype. RBC synapse with M1 ipRGC proximal dendrites or soma directly to drive the PLR.

(**B**) A schematic showing an AII amacrine cell inducing disinhibition to an M1 ipRGCs. In this circuit, AII amacrine cells most likely inhibit another amacrine cell that acts on M1 ipRGCs such as the dopaminergic amacrine cell.

(**C**) A schematic showing an AII amacrine cell coupled to Type 6 ON CBCs that synapse directly ipRGCs. Type 6 ON CBCs make en passant synapses with M1 ipRGCs and conventional synapses with M2 and M4 ipRGCs. This circuit is not required to drive the rod to RBC synapse dependent photopic PLR.

**Supplementary Figure 3 (Related to Figure 1). Anatomical confirmation of non-responsive rod and cone bipolar cells in retinal slices.**

Cells in retinal slices were filled with Alexa 750 during patch-clamp recordings to confirm their identity. The corresponding recordings of the filled cells are shown to the right. The flash intensities were the same as used in Figure 1 of the main text.

(**A**) In an Elfn1; Cx36 dKO mouse, a RBC stratifies in the ON layer of the inner plexiform layer (IPL) near the ganglion cell layer. The dashed line delineates the ON and OFF layer of the IPL. Light induced responses were absent in this cell.

(**B**) An ON CBC recorded in an Elfn1; Cx36 dKO mouse stratifies in the ON layer of the IPL. Light responses were absent in this cell.

(**C**) An OFF CBC stratifies in the OFF layer of the IPL in a mGluR6 KO mouse. The OFF CBC hyperpolarizes in response to scotopic light flashes.

**Supplementary Table 1**. **Response properties of rods and rod bipolar cells.**

Values are given as mean ± SEM, number of cells (N). The sensitivity (I_1/2_) was derived from the 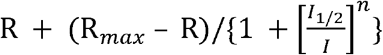. Max current is the peak of the light evoked current response.

## Notes

### Competing Interest Statement

The authors have declared no competing interest.

