## Supplementary material for "Retinal circuits driving a non-image forming visual behavior": Supp Figures 1-3

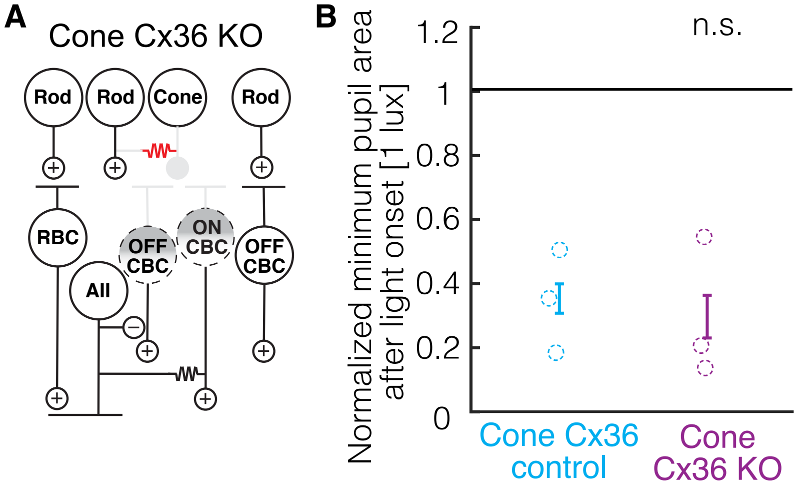


**Supplementary Figure 1. The secondary rod pathway is not required for the scotopic pupillary light response**

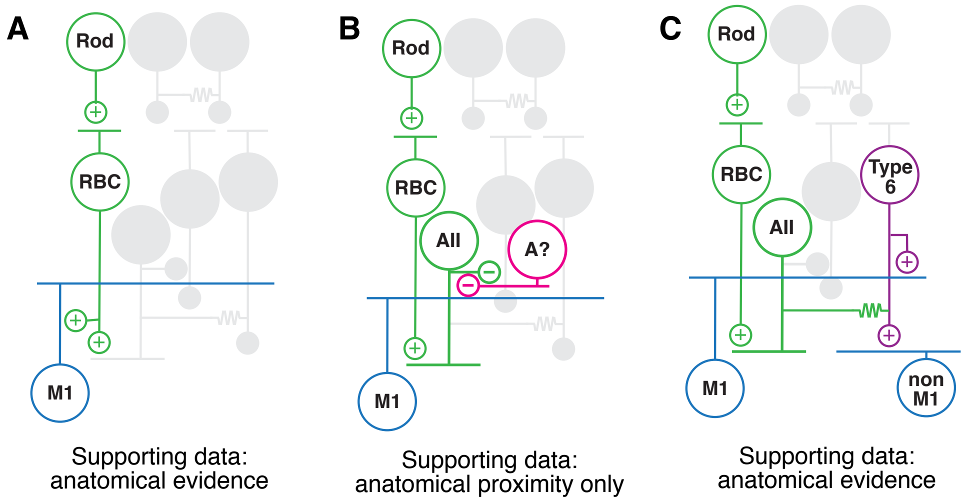


**Supplementary Figure 2. Possible rod bipolar cell to ipRGC circuits driving the PLR.**

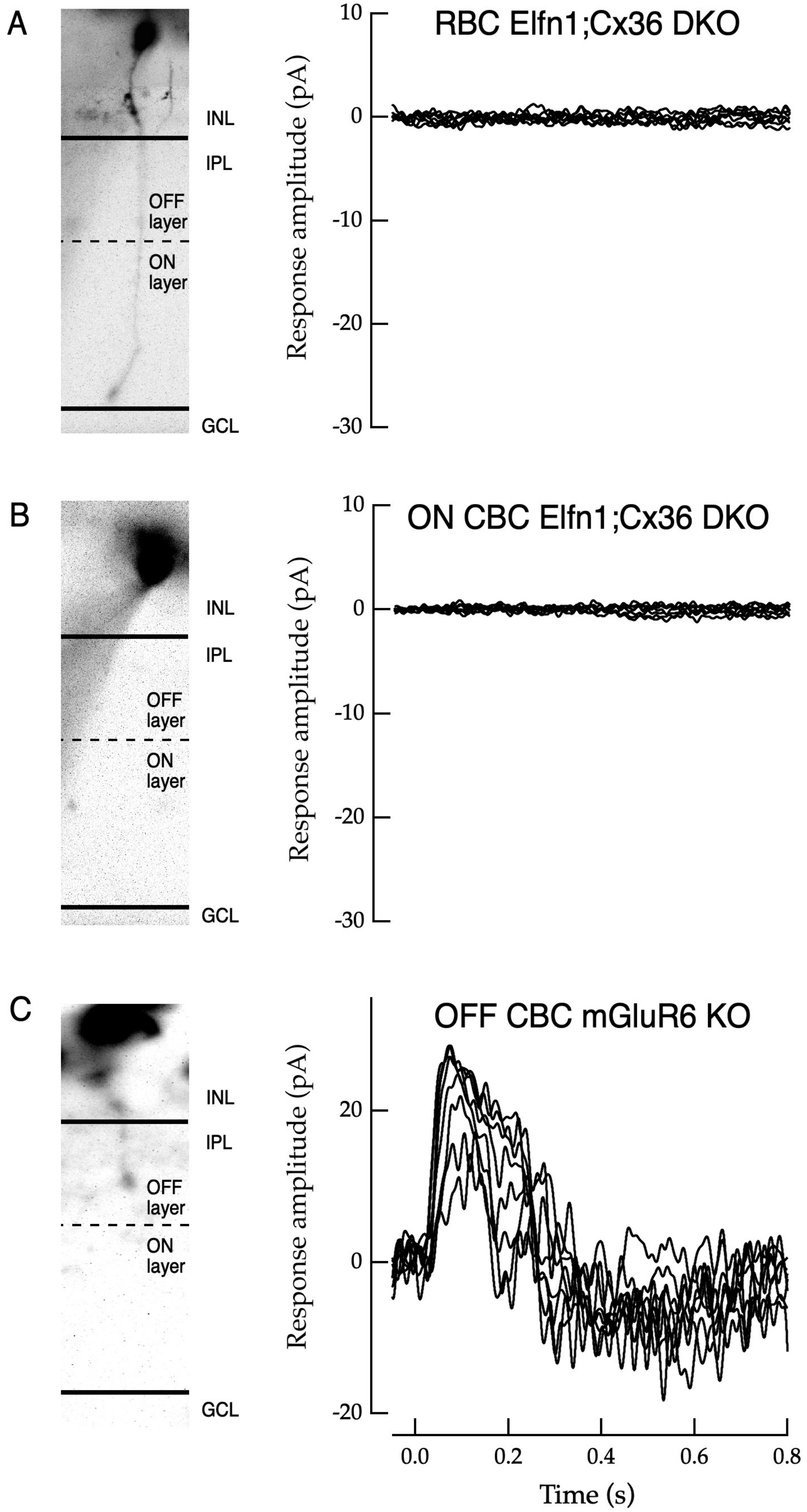


**Supplementary Figure 3 (Related to Figure 1). Anatomical confirmation of non-responsive rod and cone bipolar cells in retinal slices.**
