## Supplementary material for "Retinal circuits driving a non-image forming visual behavior": Supp Table 1

| | Sensitivity (R*/rod) | | | | MAX current (pA) | | | |
|---|---|---|---|---|---|---|---|---|
| | ROD | RBC | ON CBC | OFF CBC | ROD | RBC | ON CBC | OFF CBC |
| Control | 16.9 ± 0.7 (4) | 2.8 ± 0.2 (15) | 7.2 ± 2.2 (8) | 12.8 ± 2.2 (7) | 25.6 ± 6.7 (4) | 179.2 ± 21.6 (15) | 15.8 ± 5.5 (9) | 21.5 ± 4.6 (6) |
| mGluR6 -/- | 16.7 ± 1.2 (6) | - (4) | - (5) | 8.8 ± 1.8 (4) | 21.6 ± 4 (5) | - (4) | - (5) | 14.7 ± 5 (4) |
| Elfn1 -/- | 14.5 ± 1.3 (6) | - (10) | 6.4 ± 1.3 (10) | 9.9 ± 0.2 (9) | 27.4 ± 2.1 (7) | - (8) | 13.2 ± 2.5 (10) | 22 ± 3.9 (8) |
| Cx36 -/- | 19.8 ± 1.4 (4) | 2.6 ± 0.2 (8) | - (5) | 7.1 ± 1 (4) | 21.5 ± 3.6 (5) | 115.6 ± 24.4 (8) | - (5) | 6.4 ± 1 (5) |
| Elfn1 -/-; Cx36 -/- | 15.2 ± 1.1 (2) | - (4) | - (4) | 12.5 ± 1.3 (3) | 17.3 ± 0.5 (2) | - (4) | - (4) | 6.3 ± 1.2 (3) |

**Supplementary Table 1. Response properties of rods and rod bipolar cells.**
